# α-Synuclein plasma membrane localization correlates with cellular phosphatidylinositol polyphosphate levels

**DOI:** 10.1101/2020.07.31.224600

**Authors:** Cédric Eichmann, Reeba Susan Jacob, Alessandro Dema, Davide Mercadante, Philipp Selenko

## Abstract

The Parkinson’s disease protein α-synuclein (αSyn) promotes membrane fusion and fission by interacting with various negatively charged phospholipids. Despite postulated roles in endocytosis and exocytosis, plasma membrane (PM) interactions of αSyn are poorly understood. Here, we show that phosphatidylinositol 4,5-bisphosphate (PIP_2_) and phosphatidylinositol 3,4,5-trisphosphate (PIP_3_), two highly acidic components of inner PM leaflets, mediate plasma membrane localization of endogenous pools of αSyn in A2780, HeLa, SH-SY5Y and SK-MEL-2 cells. We demonstrate that αSyn binds reconstituted PIP_2_-membranes in a helical conformation *in vitro* and that PIP_2_ synthesizing kinases and hydrolyzing phosphatases reversibly redistribute αSyn in cells. We further delineate that αSyn-PM targeting follows phosphoinositide-3 kinase (PI3K)-dependent changes of cellular PIP_2_ and PIP_3_ levels, which collectively suggests that phosphatidylinositol polyphosphates contribute to αSyn’s cellular function(s) at the plasma membrane.

## Introduction

Aggregates of human α-synuclein (αSyn) constitute the main components of Lewy body inclusions in Parkinson’s disease (PD) and other synucleinopathies^1^. αSyn is expressed throughout the brain and abundantly found in presynaptic terminals of dopaminergic neurons, where it is involved in synaptic vesicle clustering and trafficking^2^. Whereas isolated αSyn is disordered in solution, residues 1-100 adopt extended or kinked helical conformations upon binding to membranes containing negatively charged phospholipids^3^. Complementary electrostatic interactions between lysine residues within αSyn’s N-terminal KTKEGV-repeats and acidic phospholipid headgroups align these α-helices on respective membrane surfaces^4^. Membrane curvature^5^, lipid packing defects^6, 7^and fatty acid compositions^8, 9^act as additional determinants for membrane binding. αSyn remodels target membranes^10, 11^, which likely relates to its biological function(s) in vesicle docking, fusion and fission^2^. Furthermore, αSyn multimerization and aggregation may initiate at membrane surfaces, which holds important ramifications for possible cellular scenarios in PD^9^. Early αSyn oligomers bind to and disrupt cellular and reconstituted membranes^12, 13^, whereas mature aggregates are found closely associated with membranous cell structures and intact organelles in cellular models of Lewy body inclusions^14^ and in post-mortem brain sections of PD patients^15^.

Phosphatidylinositol phosphates (PIPs) are integral components of cell membranes and a universal class of acidic phospholipids with key functions in biology^16^. Reversible phosphorylation of their inositol headgroups at positions 3, 4 and 5 generates seven types of PIPs, which act as selective binding sites for folded and disordered PIP-interaction domains^17^. In eukaryotic cells, PIPs make up less than 2% of total phospholipids with phosphatidylinositol 4,5-bisphosphate, PI(4,5)P_2_, or PIP_2_ hereafter, as the most common species^18^. PIPs function as core determinants of organelle identity^19^. PIP_2_ is predominantly found at the inner leaflet of the plasma membrane (PM), where it acts as a signaling scaffold and protein-recruitment platform^20^. Carrying a negative net charge of −4 at pH 7 renders it more acidic than other cellular phospholipids such as phosphatidylserine (net charge −1) or phosphatidic acid (net charge −1)^21^. Disordered PIP_2_-binding domains contain stretches of polybasic residues that establish complementary electrostatic contacts with the negatively-charged phosphatidylinositol-phosphate head-groups^18^ reminiscent of how αSyn KTKEGV-lysines interact with acidic phospholipids^22^. Indeed, αSyn has been shown to bind to reconstituted PIP_2_ vesicles *in vitro*^23^. Phosphatidylinositol 3,4,5-trisphosphate, PI(3,4,5)P_3_, or PIP_3_ hereafter, harbors an additional phosphate group, which renders it even more acidic (net charge −5 at pH 7)^21^. The steady-state abundance of PIP_3_ at the PM is low^16^ but local levels increase dynamically in response to cell signaling, especially following phosphatidylinositol-3 kinase (PI3K) activation^24^.

Here, we set out to investigate whether native αSyn interacted with plasma membrane PIP_2_ and PIP_3_ in mammalian cells. Using confocal and total internal reflection fluorescence microscopy, we show that endogenous αSyn forms discrete foci at the PM of human A2780, HeLa, SH-SY5Y and SK-MEL-2 cells and that the abundance and localization of these foci correlate with pools of PM PIP_2_ and PIP_3_. We further delineate high-resolution insights into αSyn interactions with reconstituted PIP_2_ vesicles by nuclear magnetic resonance (NMR) spectroscopy and establish that αSyn binds PIP_2_ membranes in its characteristic helical conformation.

## Results

### PM localization of endogenous αSyn

To determine the intracellular localization of αSyn, we selected a panel of human cell lines (A2780, HeLa, SH-SY5Y and SK-MEL-2) that expressed low but detectable amounts of the endogenous protein. Confocal immunofluorescence localization in A2780 cells with an antibody that specifically recognizes αSyn without cross-reacting with its β- and γ-isoforms (**Figure 1 – figure supplement 1A**), revealed a speckled intracellular distribution with distinct αSyn foci at apical and basal PM regions (**Figure 1A**). We verified overall antibody specificity by downregulating αSyn expression via siRNA-mediated gene silencing, which established that αSyn foci corresponded to endogenous protein pools (**Figure 1B** and **Figure 1 – figure supplement 1B**). To investigate colocalization of αSyn with PM PIP_2_, we co-stained A2780 cells with antibodies against αSyn and PIP_2_ (**Figure 1C**). In 10-20% of cases, we detected clear superpositions of αSyn and PIP_2_ signals, which we confirmed by measuring fluorescence intensity profiles over individual cell cross-sections (**Figure 1D**). To test whether changes in cellular PIP_2_ levels affected αSyn abundance at the PM, we transiently over-expressed green fluorescent protein (GFP)-tagged phosphatidylinositol-4-phosphate 5-kinase PIPKIγ^25^. PIPKIγ localizes to the PM via a unique di-lysine motif in its activation-loop^26^. Upon kinase expression, confirmed by GFP fluorescence, we detected increased amounts of αSyn at the PM of transfected cells (**Figure 1E**). By contrast, expression of GFP alone did not alter αSyn levels. We obtained similar results in HeLa and SH-SY5Y transfected cells (**Figure 1E** and **Figure 1 – figure supplement 1C**). These findings suggested that PM localization of endogenous αSyn correlated with cellular PIP_2_ levels. To better resolve the presence of αSyn at the PM, we resorted to total internal reflection fluorescence (TIRF) microscopy. Employing a narrow evanescent field depth of ~50 nm, we detected endogenous αSyn at PM foci in A2780, HeLa, SH-SY5Y and SK-MEL-2 cells, which correlated with the abundance of total αSyn determined by semi-quantitative Western blotting (**Figure 1 – figure supplement 1D**).

**Figure 1:**
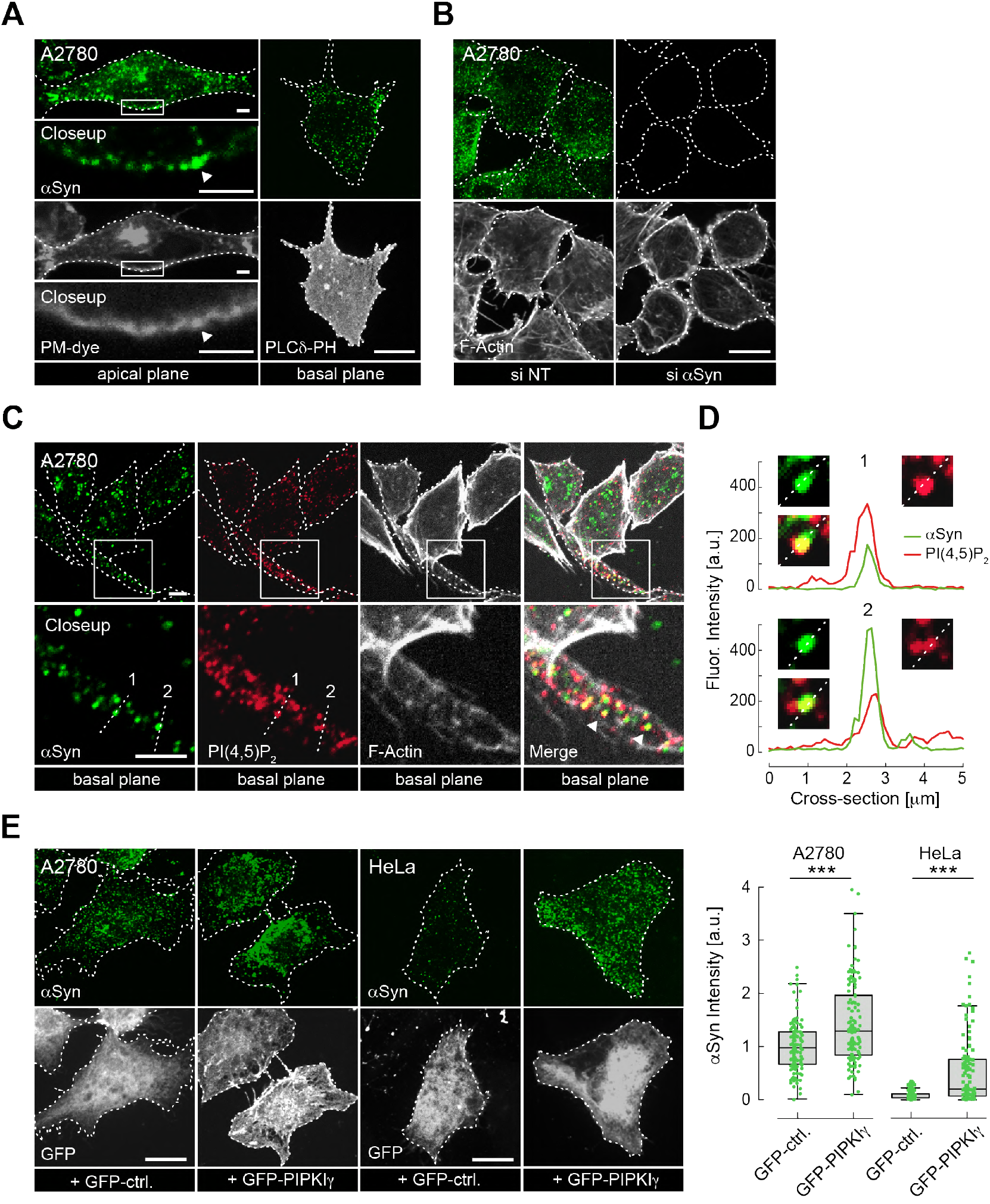
PM localization of endogenous αSyn. **(A)** Immunofluorescence detection of endogenous αSyn in A2780 cells by confocal microscopy. Plasma membranes (PM) stained with tetramethylrhodamine-WGA (left panel) or identified via PLCδ-PH-GFP (right panel). Representative apical and basal confocal planes are shown. Scale bars are 2 μm (left) and 10 μm (right). **(B)** αSyn-PM localization in A2780 cells following control (si NT) and targeted siRNA (si αSyn) knockdown. Phalloidin staining of F-Actin marks cell boundaries. Scale bars are 10 μm. **(C)** Immunofluorescence detection of endogenous αSyn and PIP_2_ at the PM. Scale bars are 5 μm. **(D)** Spatially resolved αSyn (green) and PIP_2_ (red) fluorescence intensity profiles across the dotted lines in the closeup views of (C). Resolved αSyn and PIP_2_ traces are marked with arrowheads in (C). **(E)** αSyn-PM localization and quantification after transient GFP or GFP-PIPKIγ overexpression in A2780 and Hela cells. GFP-fluorescence identifies transfected cells. Scale bars are 10 μm. Box plots for αSyn immunofluorescence quantification. Data points represent n = ~120 cells collected in four independent replicate experiments. Box dimensions represent the 25^th^ and 75^th^ percentiles, whiskers extend to the 5^th^ and 95^th^ percentiles. Data points beyond these values were considered outliers. Significance based on Student’s *t* tests as (***) P < 0.001.

### αSyn binds reconstituted PIP_2_ vesicles

To test whether αSyn directly bound PIP_2_ membranes under physiological salt and pH conditions (150 mM, pH 7.0), we added N-terminally acetylated, ^15^N isotope-labeled αSyn to reconstituted PIP_2_ vesicles. Circular dichroism (CD) spectroscopy revealed characteristic helical signatures^27, 28^ (**Figure 2A**), whereas NMR experiments confirmed site-selective line-broadening of N-terminal residues 1-100, confirming membrane binding^29, 30^ (**Figure 2B** and **Figure 2 – figure supplement 1**). In line with these observations, we detected remodeled PIP_2_ vesicles by negative-stain transmission electron microscopy (EM), manifested by tubular extrusions emanating from reconstituted specimens and agreeing with published findings on other membrane systems^10, 11^ (**Figure 2A**). Together, these results established that residues 1-100 of αSyn interacted with PIP_2_ vesicles in helical conformations that imposed membrane remodeling. To gain further insights into αSyn-PIP_2_ interactions, we reconstituted phosphatidylcholine (PC):PIP_2_ vesicles (~100 nm diameter) at fixed molar ratios of 9:1 (**Figure 2C**). We added increasing amounts of these PC-PIP_2_ vesicles to αSyn and measured CD and dynamic light scattering (DLS) spectra of the resulting mixtures. Up to a 50-fold molar excess of lipid to protein, αSyn interacted with PC-PIP_2_ vesicles in a helical conformation without disrupting the monodisperse nature of the specimens, i.e. without membrane remodeling (**Figure 2C** and **Figure 2 – figure supplement 2A**). In parallel, we performed NMR experiments on these samples and measured intensity changes of αSyn resonances in a residue-resolved manner (**Figure 2D** and **Figure 2 – figure supplement 2B**). Analyzing signal ratios (I/I_0_) of unbound and PC-PIP_2_-bound αSyn, we found that residues 1-10 constituted the primary interaction sites, whereas residues 10-100 displayed progressively weaker membrane contacts. In agreement with our experiments on PIP_2_-only vesicles, we detected no contributions by C-terminal αSyn residues. These findings confirmed the tri-segmental nature of αSyn-PIP_2_ interactions and the importance of anchoring contacts by N-terminal αSyn residues, similar to other membrane systems^29, 31, 32^.

**Figure 2:**
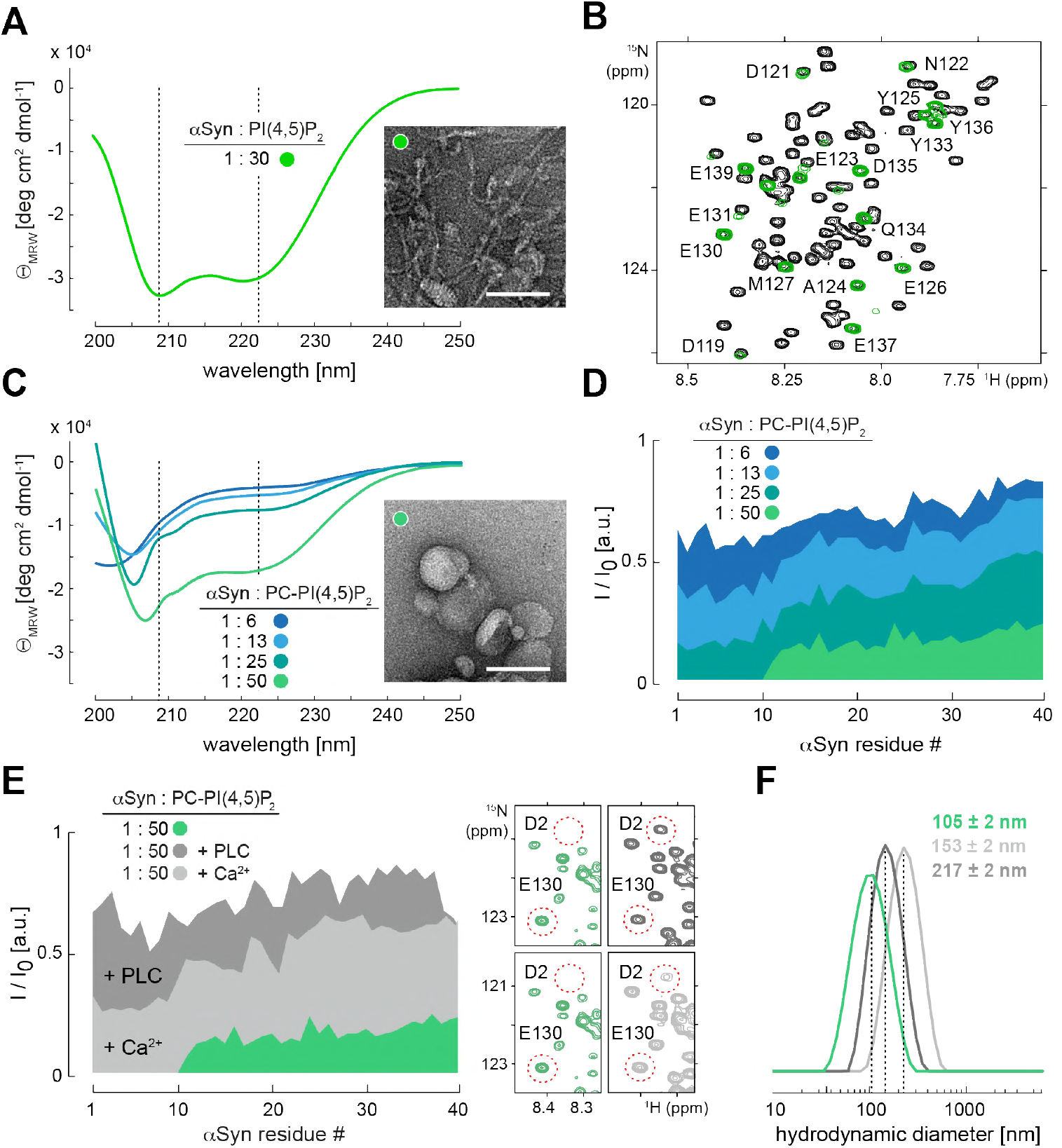
αSyn binding to reconstituted PIP_2_-vesicles. (**A**) Circular dichroism (CD) spectrum and negative-stain electron micrograph of αSyn-bound PIP_2_ vesicles (100 %). Scale bar is 100 nm. (**B**) Overlay of 2D ^1^H-^15^N NMR spectra of isolated αSyn in buffer (black) and bound to PIP_2_ vesicles (green). Remaining signals of C-terminal αSyn residues are labeled. (**C**) CD spectra of αSyn bound to PC-PIP_2_ vesicles at increasing lipid-to-protein ratios (inset) and negative-stain electron micrograph of the αSyn:PC-PIP_2_ (1:50 protein:PIP_2_) sample. Scale bar is 100 nm. (**D**) NMR signal intensity blots of bound (I) over unbound (I_0_) αSyn in the presence of different amounts of PC-PIP_2_ vesicles (equivalent to (C)). Only residues 1-40 are shown. (**E**) I/I_0_ of free (I_0_) versus PC-PIP_2_ bound αSyn at 1:50 (green, I) and after addition of PLC (dark grey) and Ca^2+^ (light grey). Selected region of 2D ^1^H-^15^N NMR spectra of PC-PIP_2_ bound αSyn (left), and in presence of PLC and Ca^2+^ (right). Vesicle release of N-terminal αSyn residues and reappearance of corresponding NMR signals are indicated for D2 (exemplary). (**F**) Hydrodynamic diameters of αSyn-bound PC-PIP_2_ vesicles before (green) and after PLC (dark grey) and Ca^2+^ (light grey) addition by dynamic light scattering (DLS). Errors were calculated based on measurements on three independent replicate samples.

To further validate our conclusions, we performed NMR experiments with mutant forms of αSyn in which we deleted residues 1-4 (ΔN)^33^, substituted Phe4 and Tyr39 with alanine (F4A-Y39A)^34^, or oxidized αSyn Met1, Met5, Met116 and Met123 to methionine-sulfoxides (MetOx)^35^ (**Figure 2 – figure supplement 3A**). In line with earlier reports, we did not observe binding to PC-PIP_2_ vesicles for any of these variants. Our results corroborated that PC-PIP_2_ interactions strongly depended on intact N-terminal αSyn residues, with critical contributions by Phe4 and Tyr39, and requiring Met1 and Met5 in their reduced states.

In contrast to other lipids, phosphatidylinositol phosphates offer attractive means to regulate the reversibility of αSyn-membrane interactions. Different charge states of PIPs can be generated from phosphatidylinositol (PI) precursors by action of PIP kinases and phosphatases^36^ or via PIP conversion by lipases such as phospholipase C (PLC) to produce soluble inositol 1,4,5-trisphosphate (IP_3_) and diacylglycerol (DAG)^37^ (**Figure 2 – figure supplement 3B**). To investigate the reversibility of αSyn-PIP_2_ interactions, we prepared PC-PIP_2_ vesicles bound to ^15^N isotope-labeled αSyn to which we added catalytic amounts of unlabeled PLC. We reasoned that PLC will progressively hydrolyze PIP_2_ binding sites and, concomitantly, release αSyn. In turn, we expected to observe an increase of αSyn NMR signals corresponding to the fraction of accumulating, unbound protein molecules. Indeed, we detected the recovery of αSyn NMR signals upon PLC addition (**Figure 2E** and **Figure 2 – figure supplement 3C**).

Next, we asked whether αSyn binding to PC-PIP_2_ vesicles was sensitive to calcium, a competitive inhibitor of many protein-PIP_2_ interactions^38^. Whereas overall binding was greatly reduced, we found that the first ten residues of αSyn displayed residual anchoring contacts with PC-PIP_2_ vesicles even at high (2.5 mM) calcium concentrations (**Figure 2E** and **Figure 2 – figure supplement 4A**), confirming earlier results on the stability of αSyn PC-PIP_2_ vesicle interactions in the presence of calcium^23^. Notably, DLS measurements showed that hydrodynamic diameters of PC-PIP_2_ vesicles expanded upon PLC treatment and in the presence of calcium, irrespective of whether αSyn was bound (**Figure 2F** and **Figure 2 – figure supplement 4B**). This further suggested that vesicle remodeling and concomitant curvature reductions did not abolish αSyn interactions. Finally, we sought to determine whether electrostatic interactions with acidic PIP headgroups alone mediated αSyn binding. To this end, we added a 4-fold molar excess of free inositol polyphosphate (IP_6_) to ^15^N isotope-labeled αSyn. Surprisingly, we did not detect binding of αSyn to this highly negatively-charged entity (**Figure 2 – figure supplement 4C**), which insinuated that αSyn interactions with PIP-containing membranes required additional lipid contributions besides headgroup contacts.

### αSyn-PM localization correlates with changes in PIP_2_-PIP_3_ levels

Following these results, we asked whether reversible αSyn-PIP_2_ interactions were present in cells. To answer this question, we transiently overexpressed different PM-targeted PIP phosphatases in A2780 cells and quantified PM localization of endogenous αSyn by confocal immunofluorescence microscopy (**Figure 3A**). Specifically, we expressed MTM1-mCherry-CAAX, which targets PI(3)P to yield phosphatidylinositol (PI), INPP5E-mCherry-CAAX to produce PI(4)P from PIP_2_, and PTEN-mCherry-CAAX to create PIP_2_ from PI(3,4,5)P_3_, as described^39^. In agreement with our hypothesis, only the conversion of PIP_2_ to PI(4)P by INPP5E led to a marked reduction of endogenous αSyn at the PM (**Figure 3A**). Together with earlier kinase results, these findings corroborated that PM localization of cellular αSyn was modulated by PIP_2_-specific enzymes. Next, we asked whether signaling-dependent activation of phosphoinositide 3-kinase (PI3K) and concomitant accumulations of the even more negatively-charged phosphatidylinositol 3,4,5-trisphosphate (PIP_3_)^24^ led to dynamic changes of αSyn abundance at the PM. To this end, we employed histamine stimulation of SK-MEL-2 cells that we transiently co-transfected with histamine 1 receptor 1 (H1R) and a PH-domain GFP-fusion construct of the general receptor of phosphoinositides 1 (GRP1) that specifically interacts with cellular PIP_3_^40^. Because histamine-mediated PI3K activation also induces time-dependent secondary effects including PIP_2_ hydrolysis by PLC^41^, we monitored αSyn localization and PIP_2_-PIP_3_ levels in a time-resolved fashion by fixing SK-MEL-2 cells at 40, 85, 120 and 240 s after histamine addition (**Figure 3B**). After 40 s, we observed an initial increase of PIP_2_ and PIP_3_ levels at the PM, which was mirrored by greater pools of endogenous αSyn at basal membrane regions. While PIP_2_ levels dropped at intermediate time-points (40-120 s), likely due to PLC-mediated PIP_2_ hydrolysis, PIP_3_ concentrations were highest at 85 s and leveled off more slowly (120-240 s). Interestingly, PM-αSyn followed the observed PIP_3_ behavior in a remarkable similar manner. At later time points (240 s), we noted a significant redistribution of cellular PIP_2_ and PIP_3_ pools towards the edges of SK-MEL-2 cells, coinciding with the accumulation of bundled Actin fibers and in line with expected PI3K-signaling-dependent rearrangements of the cytoskeleton^24^. Strikingly, αSyn colocalization with these peripheral PIP_2_-PIP_3_ speckles was significantly higher than at earlier time-points (**Figure 3B** and **Figure 3 – figure supplement 1A**). We independently confirmed these results with single time-point measurements by TIRF microscopy (**Figure 3 – figure supplement 1B**). To investigate whether other PI3K pathways caused similar effects, we stimulated SK-MEL-2 with insulin, which triggers PI3K activation via receptor tyrosine kinase (RTK) signaling^42^. We verified that SK-MEL-2 cells endogenously expressed the insulin-like growth factor receptor 1β (IGFR-1β) by Western blotting (**Figure 3 – figure supplement 1C**). In support of our hypothesis, we measured increased αSyn-PM localization by TIRF microscopy upon insulin stimulation for 10 min (**Figure 3 – figure supplement 1D**). Given the short exposure times to histamine and insulin in these experiments, we reasoned that observed PM accumulations likely reflected enhanced recruitment of existing αSyn pools rather than *de novo* protein synthesis and PM targeting, thus providing further evidence that αSyn abundance at the PM correlated with signaling-dependent changes of PIP_2_ and PIP_3_ levels.

**Figure 3:**
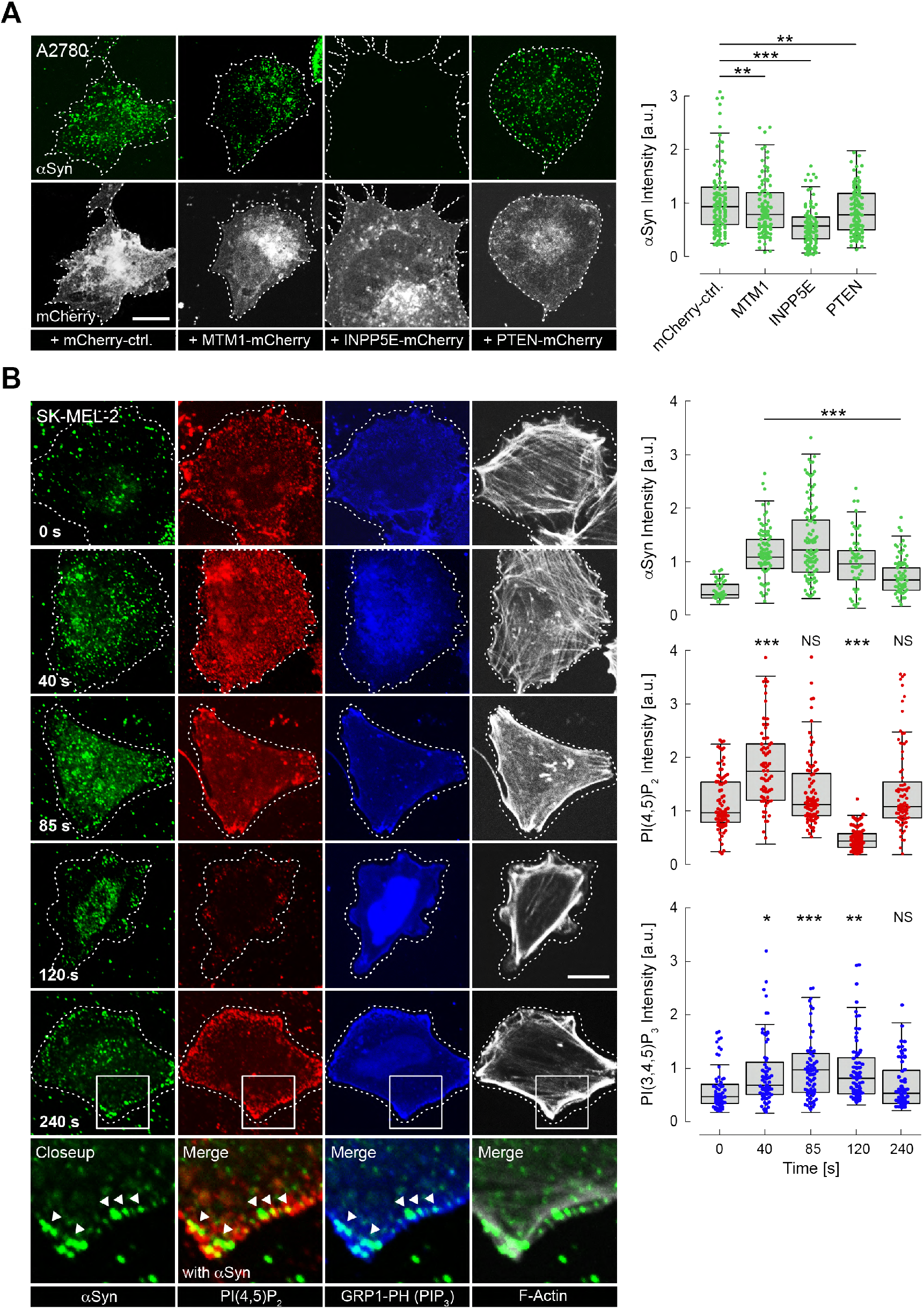
Reversible αSyn-PM localization. **(A)** Representative immunofluorescence localization of αSyn at basal A2780 PM planes by confocal microscopy. Cells transiently express different PM-targeted, mCherry-tagged PIP phosphatases, with mCherry fluorescence indicating successful transfection and phosphatase expression. A phosphatase-inactivated null mutant serves as the negative control (left). Box plots of αSyn immunofluorescence quantification are shown on the right. ~120 data points were collected per cell (n=120) in four independent replicate experiments. Box dimensions represent the 25^th^ and 75^th^ percentiles, whiskers extend to the 5^th^ and 95^th^ percentiles. Data points beyond these values were considered outliers. Significance based on ANOVA tests with Bonferroni’s post-tests as (∗∗)P < 0.01; (∗∗∗)P < 0.001. **(B)** Time-course experiments following histamine stimulation of SK-MEL-2 cells transiently expressing hH1R and GRP1-PH pEGFP-C1. Immunofluorescence detection of endogenous PIP_2_ and αSyn by confocal microscopy of basal PM regions. GRP1-PH GFP-signals report on the presence of PIP_3_. Phalloidin staining of F-Actin marks cell boundaries. Scale bar is 10 μm. Box plots represent data points collected per cell (n=80) from a single experiment, but representative of three independent experiments with similar results. Significance based on Student’s *t* tests as (∗∗)P < 0.01; (∗∗∗)P < 0.001.

## Discussion

Our results establish that clusters of endogenous αSyn are found at the plasma membrane of human A2780, HeLa, SH-SY5Y and SK-MEL-2 cells, where their native abundance correlates with PIP_2_ levels (**Figure 1**). Specifically, we show that targeted overexpression of the PIP_2_-generating kinase PIPKIγ increases endogenous αSyn at the PM (**Figure 1C**), whereas the PIP_2_-specific phosphatase INPP5E reduces the amount of PM αSyn (**Figure 3A**). We further demonstrate that PIP_3_-dependent histamine and insulin signaling redistributes αSyn to the PM (**Figure 3B** and **Figure 3 – figure supplement 1**), which collectively suggests that changes in PM PIP_2_ and PIP_3_ levels affect intracellular αSyn localization in a dynamic and reversible manner. Aiming for a stringent analysis, we investigated PM interactions at native αSyn expression levels and in a strictly unaltered sequence context, i.e., without modifying the protein with fluorescent dyes or fusion moieties. These requirements precluded live-cell imaging experiments to determine PM-localization kinetics, although histamine and insulin experiments suggest that endogenous αSyn pools redistribute readily. Although we cannot rule out that additional secondary protein-protein interactions contribute to PM targeting, we demonstrate that αSyn directly interacts with reconstituted PIP_2_ vesicles *in vitro* (**Figure 2A-D**). Importantly, the biophysical characteristics of these interactions are indistinguishable from other previously identified, negatively charged membrane systems^29, 31, 43^. Based on the known membrane-binding preferences of αSyn, PIP_2_ and PIP_3_ lipids constitute intuitive ligands. Not only because of their highly acidic nature^21^, but also because of the compositions of their acyl chains, containing saturated stearic-(18:0) and polyunsaturated arachidonic-acids (20:4), the latter conferring ‘shallow’ membrane defects^44^ ideally suited to accommodate αSyn’s helical conformations^7, 45^. Thus, from a biophysical point of view, phosphatidylinositol polyphosphates satisfy many of the known requirements for efficient membrane binding. From a biological point of view, PIPs are ubiquitously expressed and stringently required for exocytosis and endocytosis, especially in neurons, where highly abundant PIP_2_ and PIP_3_ clusters (up to ~6 mol%) mark synaptic vesicle (SV) uptake and release sites^46^. Multiple PIP-binding proteins mediate key steps in SV transmission and recycling^47, 48^and although αSyn has been implicated in synaptic exocytosis and endocytosis, its role(s) in these processes is ill defined^49^.

A2780, HeLa, SH-SY5Y and SK-MEL-2 cells are poor surrogates for primary neurons and discussing our results in relation to possible scenarios at the synapse is futile. Endogenous levels of αSyn in the tested cell lines are low, especially in comparison to presynaptic boutons, where αSyn concentrations reach up to 50 μM^50^. Similarly, the abundance of PIP_2_ and PIP_3_ is much smaller than at presynaptic terminals^46^. Hence, αSyn-PIP scenarios in the tested cell lines and in synaptic boutons are at opposite ends of protein and lipid concentration scales. Nonetheless, we believe that key conclusions of our study are generally valid. The affinity of αSyn to PIP_2_-vesicles has been reported to be in the low μM range^51^, similar to most other reconstituted membrane systems containing negatively-charged phospholipids^5, 23, 29, 30, 31^. In comparison, average dissociation constants for canonical PIP-binding scaffolds such as PH, C2, FYVE and ENTH domains vary between μM and mM^17, 18^. By contrast, disordered polybasic PIP-binding motifs target negatively-charged membranes with much weaker affinities and in a non-discriminatory fashion based on complementary electrostatic interactions^20^. αSyn-PIP binding may define a third class of interactions that are comparable in strength to folded protein domains, but driven, to large parts, by electrostatic contacts similar to those of poly-basic motifs^9^. Based on these affinity considerations, we speculate that αSyn may successfully compete for cellular PIP_2_-PIP_3_ binding sites with other proteins, especially when their abundance is in a comparable range. For binding scenarios at presynaptic terminals, this is likely the case.

Our findings are additionally supported by recent data showing that intracellular αSyn concentrations directly influenced cellular PIP_2_ levels and that protein reduction diminished PIP_2_ abundance, whereas αSyn overexpression increased PIP_2_ synthesis and produced significantly elongated axons in primary cortical neurons^52^. Conspicuously, these effects depended on αSyn’s ability to interact with membranes and were absent in a membrane-binding deficient mutant^52^. Because plasma membrane expansions require dedicated cycles of endocytosis and exocytosis^53^, αSyn-PIP interactions may contribute to both types of processes, as has been suggested earlier^54^. PM-specific αSyn-lipid interactions were additionally confirmed by ‘unroofing’ experiments in related SK-MEL-28 cells^55^, where endogenous protein pools co-localized with members of the exocytosis machinery including the known αSyn binding partners Rab3A^56^ and synaptobrevin-2/VAMP2^57^. Two other studies implicated αSyn and αSyn-PIP_2_ interactions in clathrin assembly and clathrin-mediated endocytosis, respectively^58, 59^, which further strengthens the notion that phosphatidylinositol polyphosphates contribute to αSyn functions at the plasma membrane.

## Materials and Methods

### Mammalian Cell Lines and Growth Media

Human cells lines A2780 (Sigma, cat.# 93112519), HeLa (Sigma, cat.# 93022013), SH-SY5Y (Sigma, cat.# 94030304) and SK-MEL-2 (provided by Ronit Sharon, Hebrew University, Israel) were grown in humidified 5 % (v/v) CO_2_ incubators at 37 °C in the following media supplemented with 10% (v/v) fetal bovine serum (FBS): RPMI 1640 (A2780), low glucose DMEM (HeLa), DMEM-Ham’s F-12 (SH-SY5Y) and MEM with 1% non-essential amino acids and 2 mM glutamine (SK-MEL-2). Cells were split at 70-80% confluence with a passage number below 20 for all experiments. All cell lines were routinely confirmed to be mycoplasma free.

### Transient Cell Transfections

A2780 cells were seeded on fibronectin (Sigma) coated 25 mm cover slips in 12-well plates at a density of 3 × 10^5^ cells. Cells were transfected using Lipofectamine 3000 (Thermo) according to manufacturers’ instructions. SK-MEL-2 cells were seeded on 18 mm coverslips at a density of 2 × 10^5^ cells and transfected using TransIT-X2 (Mirus Bio) according to manufactures’ instructions. Details of plasmids used for transfection are provided in **Table 1**. 1 μg of plasmids was used in all cases. Following transfection, cells were grown for 24 h before analysis.

**Table 1.**

| Plasmid | Function | Figure | Source | Reference |
| --- | --- | --- | --- | --- |
| EGFP-PLC$\delta_1$-PH | PH domain, binds PI(4,5)P<sub>2</sub> at PM | Fig. 1A | Dr. Michael Krauss (FMP-Berlin, Germany) | 60 |
| EGFP-phosphatidylinositol 4-phosphate 5-kinase type Iγ (PIPKIγ) | PIP Kinase, creates PI(4,5)P<sub>2</sub> at PM | Fig. 1E, Fig. 1 - S1C | Dr. Volker Haucke (FMP-Berlin, Germany) | 25 |
| MTM1-mCherry-CAAX | PIP Phosphatase, acts on PI(3)P. Targeted to PM. | Fig. 3A | Dr. Volker Haucke (FMP-Berlin, Germany) | 39 |
| INPP5E-mCherry-CAAX | PIP Phosphatase, acts on PI(4,5)P<sub>2</sub>. Targeted to PM. | Fig. 3A | Dr. Volker Haucke (FMP-Berlin, Germany) | 39 |
| PTEN-mCherry-CAAX | PIP Phosphatase, acts on PI(3,4,5)P<sub>3</sub>. Targeted to PM. | Fig. 3A | Dr. Volker Haucke (FMP-Berlin, Germany) | 39 |
| INPP4A-mCherry-CAAX (mutated) | PIP Phosphatase dead mutant. Targeted to PM | Fig. 3A | Dr. Volker Haucke (FMP-Berlin, Germany) | 39 |
| H1R | human Histamine 1 receptor | Fig. 3B | Dr. Ronit Sharon (Hebrew University, Israel) | 61 |
| GRP1-PH pEGFP-C1 | PH domain binds PI(3,4,5)P<sub>3</sub> | Fig. 3B | Addgene (Plasmid #71378) | 40 |

### siRNA Knockdown Experiments

Commercial siRNA mixtures against human αSyn (Dharmacon, ON-TARGET plus human SNCA, cat.# L-002000-00-0005) and a non-targeted control (cat.# D-001810-10-05) were used. A2780 cells were seeded at a density of 6 × 10^5^ cells and transfected with 1.7 μg of the respective siRNA mixtures using Lipofectamine 3000 according to manufacturers’ instructions. After transfection, cells were grown for 48 h before analysis.

### Immunofluorescence

For immunofluorescence (IF) imaging of endogenous αSyn and expressed PIP-kinase/phosphatases, cells were washed 3 × 5 min with PBS and fixed in 4 % (w/v) paraformaldehyde (PFA) for 15 min at room temperature (RT). For plasma membrane staining with 5 μg/mL Alexa Fluor 350/tetramethylrhodamine conjugated to Wheat Germ Agglutinin (WGA) (Invitrogen), cells were fixed and washed with PBS before application for 10 min at RT. Excess dye was washed off with PBS. For antibody staining, cells were permeabilized with 0.5% Saponin in PBS for 10 min, and blocked with 5% (w/v) bovine serum albumin (BSA, Sigma) in PBS for 30 min. After blocking, cells were incubated with anti-αSyn antibody for 90 min at RT. After washing 3 × 5 min with PBS, cover slips were incubated with Alexa Fluor-tagged secondary antibody for 45 min at RT. Details of antibodies are provided in **Table 2**. Before confocal microscopy cover slips were mounted with Immu-Mount (Thermo), after 3 × 5 min PBS washes. Immunofluorescence detection of PI(4,5)P_2_ at the PM was performed according to^62^ with slight modifications. A2780 and SK-MEL-2 cells were cultured on fibronectin-coated coverslips and pre-extracted in PHEM buffer (60 mM PIPES, 25 mM HEPES, 5 mM EGTA, 1 mM MgCl_2_) to remove the majority of soluble cytoplasmic proteins. Cells were fixed with 4% PFA and 0.2% glutaraldehyde in PHEM buffer for 15 min at RT. All post-fixation steps until mounting were carried out at 4 °C. Washes were performed with ice-cold PIPES buffer (20 mM PIPES, pH 6.8, 137 mM NaCl, 2.7 mM KCl) to minimize damage to endogenous PIP moieties. Following fixation, cells were washed thrice in PIPES buffer containing 50 mM NH_4_Cl and subsequently blocked and permeabilized in PIPES buffer supplemented with 5% ‘normal goat serum’ and 0.5% Saponin for 30 min. Post blocking, cells were incubated with anti-PI(4,5)P_2_ and anti-αSyn antibodies for 60 min, washed thrice and incubated with Alexa Fluor 647 secondary antibody for 45 min. All antibodies used in this study are listed in **Table 2**. Before confocal microscopy, cover slips were mounted with Immu-Mount (Thermo) after 3 × 5 min PIPES buffer washes.

**Table 2.**
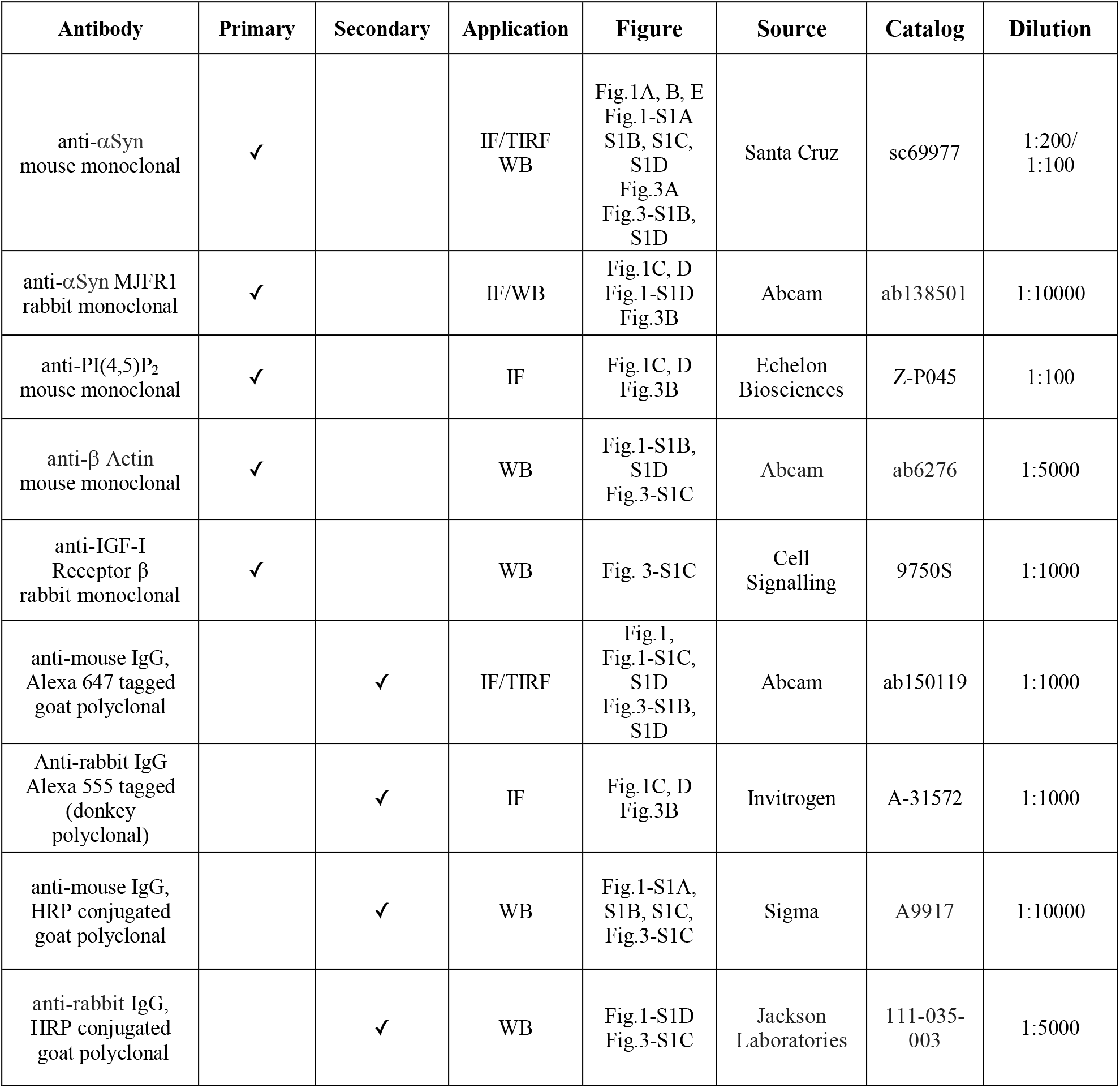

### Confocal Microscopy

Confocal microscopy imaging was performed on a Nikon Spinning Unit confocal microscope with an oil 60x objective and additional 1.5x magnification. Four channels in 5 optical sections from the basal PM plane were acquired with excitation wavelengths of 405 (blue, 50% laser power, for WGA), 488 (green, 20%, for GFP), 568 (red, 20%, for mCherry) and 647 (far-red, 20%, goat anti-mouse) with 200 ms exposure times. At least 25 images per biological replicate were collected and 3-4 replicates per experiment were analyzed

### Total Internal Reflection Fluorescence (TIRF) Microscopy

For TIRF localization of endogenous αSyn at the PM, A2780, HeLa, SH-SY5Y and SK-MEL-2 cells were cultured on 18 mm fibronectin-coated coverslips at a density of 2 × 10^5^ cells for 24 h and fixed with 4% PFA. After fixation, antibody detection was performed as described in the previous section. Coverslips for TIRF imaging were mounted in PBS after immunostaining and imaged on an Andor Dragonfly Spinning Disc microscope with a TIRF 100×/NA 1.45 oil objective. For TIRF detection of PM-proximal fluorescence signals, evanescent fields were kept at 50 nm in all experiments. Four lasers operating at 405 nm (15% laser power), 488 nm (20% laser power), 561 nm (20% laser power) and 647 nm (20% laser power) were used for fluorophore excitation, along with 200 ms exposure times for image acquisitions. At least 20 images per biological replicate were collected and 3 replicates per experiment were analyzed.

### Histamine and Insulin Stimulation

PI-3 kinase activity was stimulated either by insulin or histamine addition. For insulin stimulation via the endogenously expressed insulin-like growth factor-1 receptor (IGF-1r)^63^, SK-MEL-2 cells were seeded on coverslips and starved in HBSS for 18 h, as described^64^. 100 nM of insulin was added to cells for 10 min and cells were fixed immediately afterwards. For histamine stimulation, SK-MEL-2 cells were seeded on 18 mm coverslips at a density of 2 × 10^5^, transiently transfected with human histamine 1 receptor (hH1R) and serum-starved for 3 h, as described in^65^. 500 μM of histamine was added to cells for 40 s and cells were fixed immediately afterwards. All cell samples were further processed as previously outlined for TIRF procedures. F-Actin was detected by Phalloidin-Alexa Fluor 405 staining (1:400, Invitrogen) during secondary antibody incubation.

### Image Analysis and Quantification

Image analysis and quantification were performed in Fiji^66^. For confocal image quantification, focal planes of apical and basal PMs were selected manually. Images were segmented based on GFP signals by automatic thresholding according to Huang et al^67^. Threshold regions were marked as regions-of-interest (ROIs), copied to the far-red channel (αSyn IF) and fluorescence intensities were determined. In the box plots of **Figures 1E** and **3A, B**, each ROI corresponds to a single cell and is represented as a data point. For TIRF data in **Figure 3 – figure supplement 1B** and **1D**, images were segmented based on Phalloidin signals via automated thresholding using the default algorithm in Fiji^68^. Different than for confocal images in **Figures 1** and **3**, TIRF ROIs consist of multiple adjacent cells in a single frame that were copied to the far-red channel (αSyn IF). ROIs of less than 2 μm^2^ in size were excluded. The Fiji particle counting routine was used to determine the number of αSyn puncta in each ROI. The number of cells in each image was determined manually based on cell outlines marked by Phalloidin. In **Figure 3 – figure supplement 1B** and **1D**, data points in box plots were calculated by dividing the number of αSyn puncta per image by the cell count. All box plots depict median values (center lines) with box dimensions representing the 25^th^ and 75^th^ percentiles. Whiskers extend to 1.5-times the interquartile range and depict the 5^th^ and 95^th^ percentiles. Each box plot in **Figures 1A** and **3A** corresponds to 110-120 data points combined from three independent biological replicates. Box plots in **Figure 3B** contain data points collected per cell (n=80) from a single experiment but representative of three independent experiments with similar results. Box plots in **Figure 3 – figure supplement 1B** and **1D** contain data points from approximately 120 cells, combined from three independent biological replicates.

### Statistical Analysis

For box plots, data points considered ‘outliers’ were determined based on criteria defined in the Grubbs outlier test^69^ and omitted. ANOVA tests with Bonferroni’s post-tests^70^, ^71^ were used to determine the statistical significance of experiments with more than two samples, whereas Student’s *t* tests were performed to assess statistical differences between samples^72^. Significance is given as *P < 0.05; **P < 0.01; ***P < 0.001.

### Cell Lysate Preparation

Lysates of A2780, HeLa, SH-SY5Y, SK-MEL-2 cell lines were prepared by detaching ~5-10 million cells with trypsin/EDTA (0.05% / 0.02%) and harvested by centrifugation at 130 x g for 5 min at 25 °C. Sedimented cells were washed once with PBS, counted on a haemocytometer and pelleted again by centrifugation. After resuspending cells in PBS with proteinase inhibitor cocktail (Roche), yielding a cell count of 2 × 10^7^ cells /ml, they were lysed by repeated freeze-thaw cycles. Lysates were cleared by centrifugation at 16000 x g for 30 min. Supernatants were removed, total protein concentration measured with a BCA assay kit (Thermo) and 50 μg of protein (per lane) was applied onto SDS-PAGE for western blotting.

### Western Blotting

Cell lysates and recombinant protein samples were boiled in Laemmli buffer for 10 min before SDS-PAGE separation on commercial, precast 4-18% gradient gels (BioRad). Recombinant N-terminally acetylated α-, β- and γ-Syn, at specified concentrations were loaded as reference inputs (see Protein Expression and Purification). Proteins were transferred onto PVDF membranes and fixed with 4% (w/v) paraformaldehyde (PFA) in PBS for 1 h^73^. Membranes were washed 2x with PBS, 2x with tris-buffered saline with 0.1 % tween 20 (TBST) and blocked in 5% milk-TBST for 1 h. After blocking, the blots were incubated with primary antibodies overnight at 4 °C. Membranes were then washed and probed with HRP-conjugated secondary antibodies for 1 h. The antibodies used for each blot is provided in **Table 2**. Membranes were developed using the SuperSignal West Pico Plus reagent (Thermo) and luminescence signals were detected on a BioRad Molecular Imager.

### Western Blot Quantification

Intensities of αSyn and β-Actin bands were quantified using the ImageLab software (BioRad). αSyn reference input was used to generate a standard curve. For cell lysate samples, αSyn intensity was normalized according to the β-Actin signal and cell lysate were calculated with respect to the αSyn standard curve. Error bars denote background signal (noise).

### Recombinant Protein Expression and Purification

^15^N isotope-labeled, N-terminally acetylated, human wild-type α-, β- and γ-Syn were produced by co-expressing PT7-7 plasmids with yeast N-acetyltransferase complex B (NatB)^74^ in *Escherichia coli* BL21 Star (DE3) cells using M9 minimal media supplemented with 0.5 g/L of ^15^NH_4_Cl (Sigma). Protein purification under non-denaturing conditions was performed as described previously^75^. αSyn mutants ΔN and F4A-Y39A were generated by site-directed mutagenesis (QuikChange, Agilent) and confirmed by DNA sequencing. Recombinant protein expression and purification of αSyn F4A-Y39A was identical to wild-type αSyn. Lacking the N-terminal substrate specificity for NatB, αSyn ΔN was produced in its non-acetylated form and purified as the wild-type protein. Methionine-oxidized ^15^N isotope-labeled wild-type αSyn was expressed and purified as described^76^. Protein samples were concentrated to 1-1.2 mM in NMR buffer (25 mM sodium phosphate, 150 mM NaCl) at pH 7.0. Protein concentrations were determined spectrophotometrically by UV absorbance measurements at 280 nm with ε = 5690 M^-1^cm^-1^ for α-, β-Syn ΔN, and methionine-oxidized αSyn. For αSyn F4A-Y39A and and γ-Syn, ε = 4470 and 1490 M^-1^cm^-1^ were used. Final aliquots of protein stock solutions were snap frozen in liquid nitrogen and stored at −80 °C until use.

### Reconstituted PI(4,5)P_2_ Vesicles

Phospholipids were purchased from Avanti Polar Lipids. Small unilamellar vesicles (SUVs) were prepared from 100% brain (porcine) phosphatidylinositol 4,5-bisphosphate (PIP_2_). A thin lipid film was formed in a glass vial by gently drying 1 mg of PIP_2_ in chloroform-methanol under a stream of nitrogen. To remove residual traces of organic solvents, the lipid film was placed under vacuum overnight. 0.5 mL NMR buffer was then added to hydrate the lipid film for 1 hour at RT while agitating. After 5 freeze-thaw cycles on dry ice and incubation in a water bath at RT, the lipid suspension was sonicated at 4 °C for 20 min at 30% power setting (Bandelin). Resulting PIP_2_ SUVs (2 mg/mL) were used immediately. αSyn:PIP_2_ molar ratios for sample preparations were calculated using a PIP_2_ lipid mass of 1096 Da. For PIP_2_ titration experiments, 1, 5, 10, 15, 20, and 30-fold molar excess of lipids was added to 60 μM of ^15^N isotope-labeled, N-terminally acetylated αSyn (total volume 120 μL) and αSyn PIP_2_ samples were incubated for 45 min at RT before NMR and CD measurements. Following the same procedure, mixed phosphatidylcholine phosphatidylinositol-4,5 bisphosphate (PC:PIP_2_, 9:1) suspensions were prepared using 9 mg of 1,2-dioleoyl-sn-glycero-3-phosphocholine (DOPC, 786 Da) and 1 mg PIP_2_. The dried lipid film was hydrated with 0.25 mL NMR buffer. The PC-PIP_2_ suspension was then extruded through polycarbonate membranes with a pore size of 100 nm according to manufacturer’s instructions (mini-extruder, Avanti Polar Lipids) and resulting PC-PIP_2_ large unilamellar vesicles LUVs (40 mg/mL) were used immediately. For sample preparations, an average PC-PIP_2_ lipid mass of ~820 Da (0.9 × 786 Da + 0.1 × 1096 Da) was used to calculate the αSyn:PC-PIP_2_ molar ratios. ^15^N isotope-labeled, N-terminally acetylated αSyn (60 μM) was incubated with 80, 170, 340, and 680-fold molar excess of total PC-PIP_2_ lipids. αSyn PC-PIP_2_ samples (total volume 120 μL) were incubated for 45 min at RT before CD, NMR, and DLS experiments. Synthetic inositol hexaphosphate (IP_6_) was kindly provided by Dr. Dorothea Fiedler, Department of Chemical Biology (FMP Berlin). Before NMR measurements, 50 μM αSyn was incubated with 200 μM IP_6_ in NMR buffer (total volume 120 μL) for 45 min at RT.

### Phospholipase C Reaction

Phospholipase C (PLC) was purchased from Sigma and the lyophilized powder was dissolved in NMR buffer at 1000 units (U)/mL. αSyn PC-PIP_2_ samples at 680-fold molar excess of PC-PIP_2_ lipids (60 μM αSyn, 40 mM PC-PIP_2_) were incubated while agitating at 37 °C for 45 min with 10 U of PLC and 1 mM PMSF in a total volume of 120 μL yielding a PLC activity of ~80 mM per min.

### Nuclear Magnetic Resonance (NMR) Spectroscopy

For best comparison of protein reference and αSyn-lipid NMR data, final concentrations of ^15^N isotope-labeled, N-terminals acetylated αSyn samples were adjusted to 60 μM, supplemented with 5% D_2_O and measured in 3 mm (diameter) Shigemi tubes in all cases. NMR experiments were acquired on a Bruker 600 MHz Avance spectrometer equipped with a cryogenically cooled proton-optimized ^1^H{^13^C/^15^N} TCI probe. Reference and αSyn-lipid NMR spectra were acquired with identical spectrometer settings and general acquisition parameters. Specifically, we employed 2D ^1^H-^15^N SOFAST HMQC NMR pulse-sequences^77^ with a data size of 128 × 512 complex points for a sweep width (SW) of 28.0 ppm (^15^N) and 16.7 ppm (^1^H), 128 scans, 60 ms recycling delay, recorded at 283 K. Inspection of the highly pH-sensitive His50 ^1^H-^15^N chemical shift indicated that the sample pH changed from 7 to 6.5 during the PLC reaction (**Figure 2E**). To accurately delineate I/I_0_ values, we recorded reference NMR spectra at pH 6.5. All NMR spectra were processed with PROSA, zero-filled to four times the number of real points and processed without window function. Visualization and data analysis were carried out in CARA. NMR signal intensity ratios (I/I_0_) of isolated αSyn (I_0_) and in the presence lipids (I) were determined for each residue by extracting maximal signal peak heights in the respective 2D ^1^H-^15^N NMR spectra.

### Circular Dichroism (CD) Spectroscopy

NMR samples of isolated αSyn and αSyn in presence of lipid vesicles were diluted with NMR buffer to a final protein concentration of 10 μM for CD measurements. CD spectra (200-250 nm) were collected on a Jasco J-720 CD spectropolarimeter in a 1 mm quartz cell at 25 °C. One replicate per sample was recorded. Six scans were averaged and blank samples (without αSyn) were subtracted from the protein spectra to calculate the mean residue weight ellipticity (θ_MRW_).

### Dynamic Light Scattering (DLS)

DLS measurements were acquired on a Zetasizer Nano ZS (Malvern Instruments) operating at a laser wavelength of 633 nm equipped with a Peltier temperature controller set to 25 °C. Data were collected on all NMR samples containing αSyn, isolated PC-PIP_2_ vesicles, and PC-PIP_2_ vesicles in presence of Ca^2+^ and PLC, respectively. Using the Malvern DTS software, mean hydrodynamic diameters were calculated from three replicates of the same sample in the intensity-weighted mode.

### Negative-Stain Electron Microscopy (EM)

NMR samples of αSyn at 30- and 680-fold molar excess of PIP_2_ and PC-PIP_2_ lipids were diluted to a protein concentration of ~10 μM in NMR buffer. 5 μL aliquots were added to glow-discharged carbon-coated copper grids for 1 min. Excess liquid was removed with filter paper and grids were washed twice with H_2_O before staining with 2% (w/v) uranyl acetate for 15 s. Negative stain, transmission EM images were acquired on a Technai G2 TEM.

## Acknowledgments

We are grateful to Dr. Peter Schmieder and Monika Beerbaum for excellent maintenance of the NMR infrastructure at the Leibniz Institute of Molecular Pharmacology (FMP Berlin) and Dr. Tali Scherf for NMR infrastructure maintenance at the Weizmann Institute of Science. We thank Dr. Dmytro Puchkov (FMP Berlin) for assistance with negative-stain electron microscopy and Drs. Michael Krauss and Volker Haucke (FMP Berlin) for tools and reagents, helpful discussions throughout the project and useful feedback on the manuscript. Dr. Dorothea Fielder (FMP Berlin) for sharing aliquots of IP_6_. We also thank Drs. Martin Lehmann (Cellular imaging, FMP Berlin) and Yoseph Addadi (Life Sciences Core Facilities, Weizmann Institute of Science) for excellent maintenance of imaging facilities and their support at the respective institutes. We acknowledge highly valuable input by Drs. Meir Schechter and Ronit Sharon, Hebrew University Jerusalem, especially with regard to time-resolved histamine and insulin stimulation experiments. We further thank them for kindly providing aliquots of the SK-MEL-2 cell line. We are grateful to Drs. Ori Avinoam, Hagen Hofmann (Weizmann) and Andres Binolfi (CONICET) for carefully reading the manuscript. C.E. was supported by a Swiss National Science Foundation (SNSF) Advanced Postdoc Mobility fellowship P300PA_160979. P.S. acknowledges funding by the European Research Council (ERC) Consolidator Grant NeuroInCellNMR (647474). Work in the Selenko laboratory is supported by the Willner Family Foundation.

---

**Figure 1 - Supplement 1:**
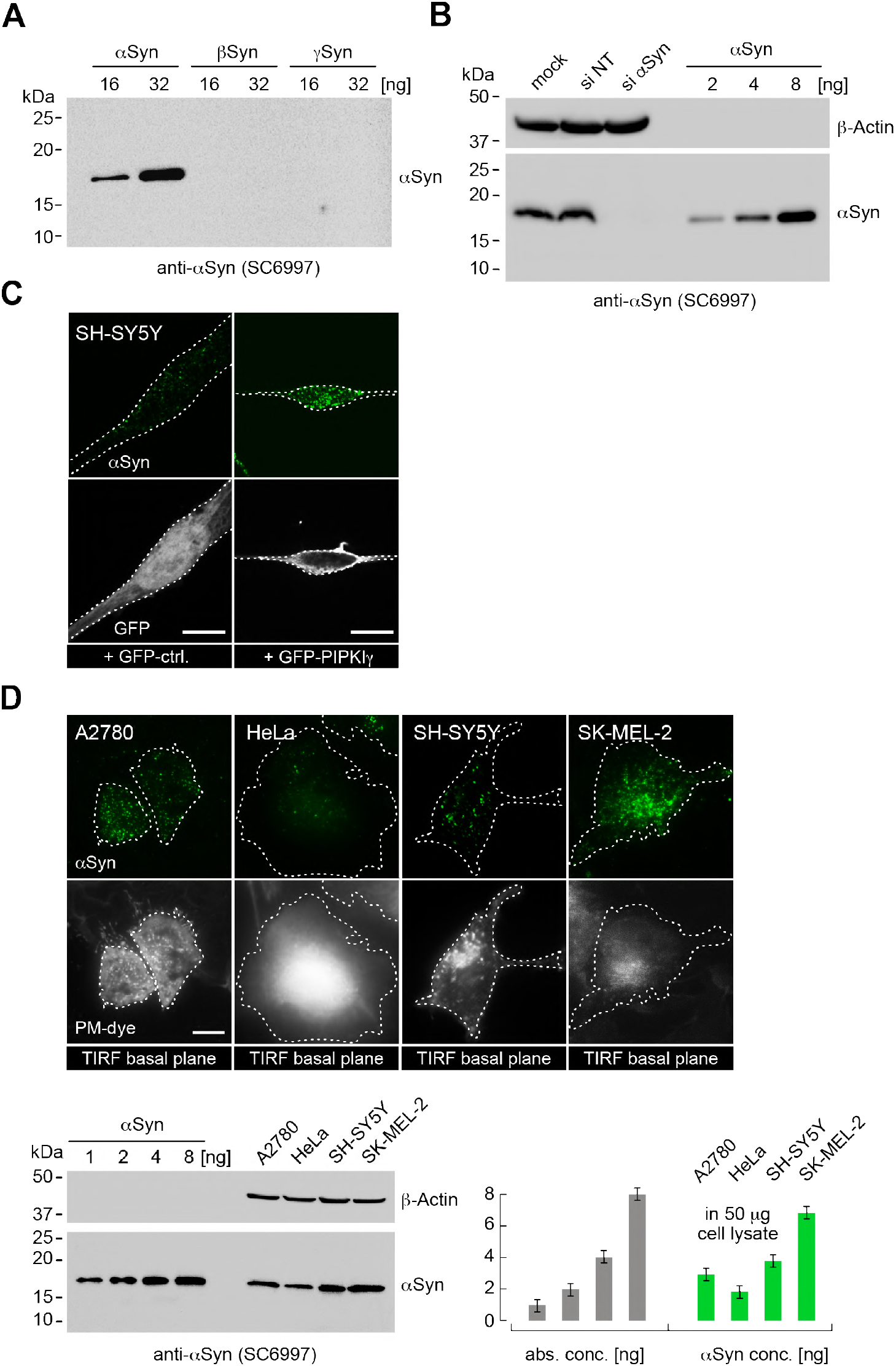
**(A)** Western Blot to determine the specificity of the αSyn antibody (sc69977) against β and γ isoforms of the protein. **(B)** Western blot of A2780 lysates of control (si NT) and targeted siRNA (si αSyn) knockdown cells. Recombinant N-terminally acetylated αSyn serves as input-, β-Actin as loading-controls. **(C)** Immunofluorescence localization of endogenous αSyn in SH-SY5Y cells transfected with GFP or PH-GFP-PIPKIγ. **(D)** Total internal reflection (TIRF) fluorescence-microscopy of αSyn-PM localization in A2780, HeLa, SH-SY5Y and SK-MEL-2 cells. PM stained with tetramethylrhodamine-WGA. Scale bars are 10 μm. Western blot of endogenous αSyn in respective cell lysates. Recombinant N-terminally acetylated αSyn serves as input-, β-Actin as loading-controls. Bar graphs denote Western blot quantifications of αSyn with signals normalized against β-Actin. Error bars denote standard deviations based on measured background signals.

**Figure 2 - Supplement 1:**
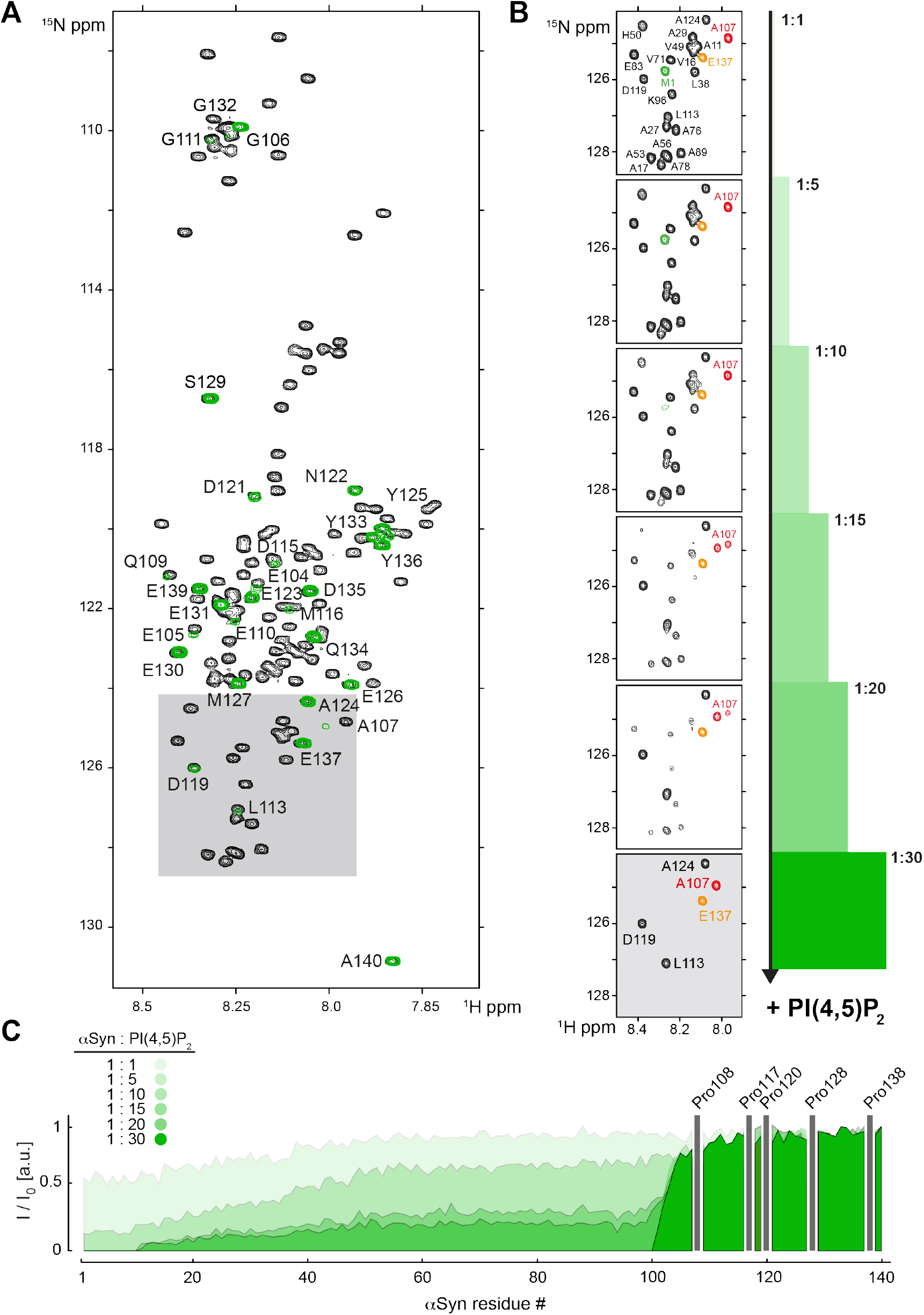
**(A)** Overlay of 2D ^1^H-^15^N NMR spectra of isolated, N-terminally acetylated αSyn (black) and bound to a 30-fold molar excess of PIP_2_-only vesicles (green). Uniform signal broadening of N-terminal residues 1-100 is evident. Observable signals of C-terminal αSyn residues 100-140 are labeled. **(B)** Selected region of 2D ^1^H-^15^N NMR spectra of αSyn upon addition of increasing amounts of PIP_2_-only vesicles, corresponding to molar protein:lipid ratios of 1:1, 1:5, 1:10, 1:15, 1:20 and 1:30, with M1 labeled in green and E137 indicated in orange. Note that A107 (red) at the border between membrane-bound (N-terminal) and - unbound (C-terminal) αSyn residues displays peak-splitting at increasing PIP_2_ concentrations, indicative of chemical shift differences between free and membrane-bound protein states. **(C)** Residue-resolved signal attenuation profiles (I/I_0_) of free (I_0_) versus PIP_2_ bound (I) αSyn, at previously indicated molar ratios. Positions of C-terminal αSyn proline residues without peptide amide resonances are shown in the three-letter amino acid code.

**Figure 2 - Supplement 2:**
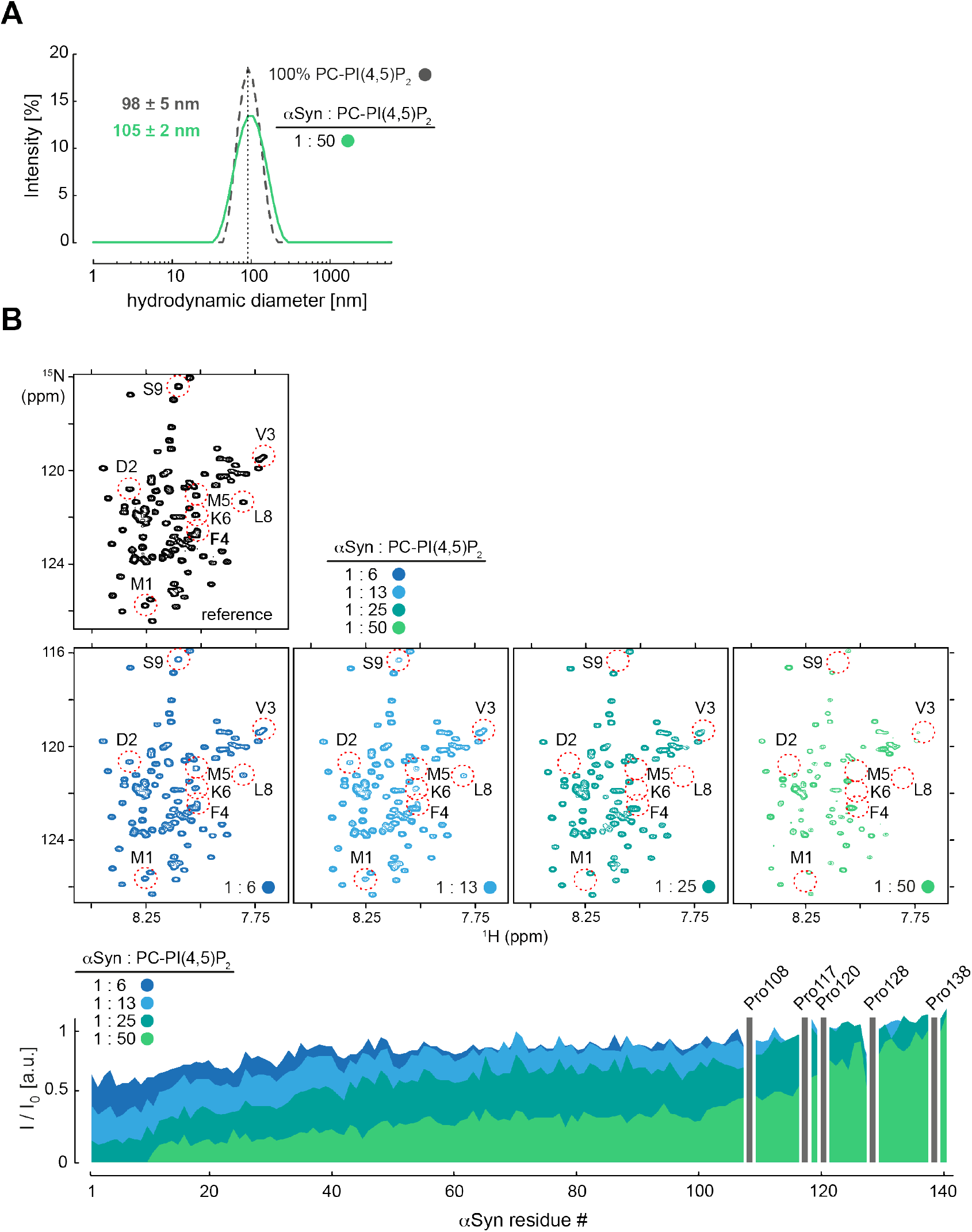
(**A**) Hydrodynamic diameters of PC-PIP_2_ vesicles in the absence (dashed grey) and presence of αSyn (green) at a protein:lipid ratio of 1:50 by DLS measurements. Errors were calculated based on measurements of three independent replicate samples. (**B**) Selected region of 2D ^1^H-^15^N NMR spectra of isolated, N-terminally acetylated αSyn (black) and in the presence of 6, 13, 25, 50 mol equivalents of PC-PIP_2_ vesicles (blue to green). Site-selective line broadening of N-terminal residues 1-10 is highlighted. Residue-resolved signal attenuation profiles (I/I_0_) of free (I_0_) versus PC-PIP_2_ bound (I) αSyn at previously indicated molar ratios. Positions of C-terminal αSyn proline residues without peptide amide resonances are shown in the three-letter amino acid code.

**Figure 2 - Supplement 3:**
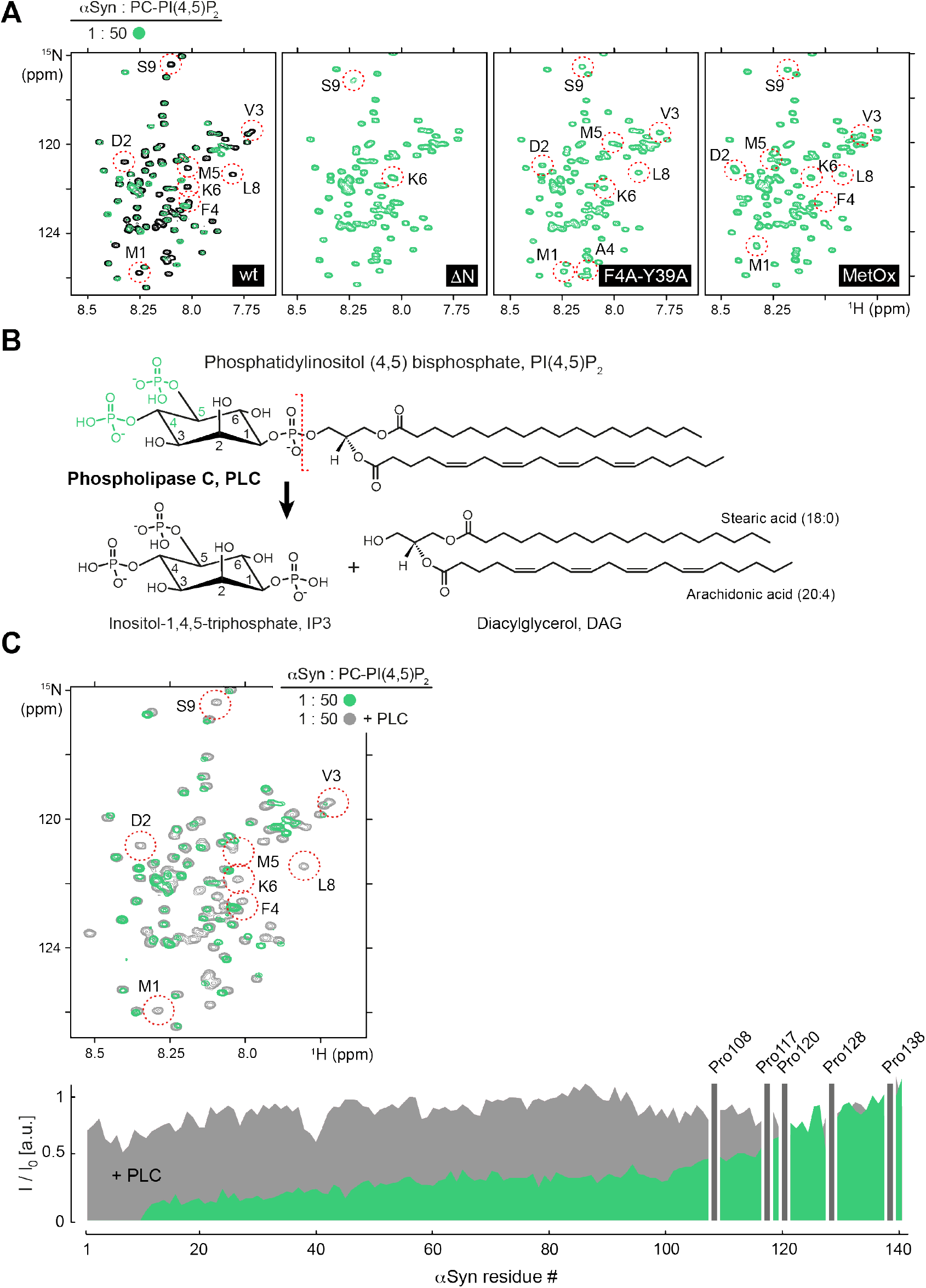
(**A**) Selected regions of 2D ^1^H-^15^N NMR spectra. Left to right: Overlay of isolated, N-terminally acetylated wild-type (WT) αSyn and bound to PC-PIP_2_ vesicles at a protein:lipid ratio of 1:50 (green). NMR spectra of N-terminally truncated αSyn lacking residues 1-5 (ΔN), mutated (F4A-Y39A) and methionine-oxidized (MetOx) αSyn in the presence of PC-PIP_2_ vesicles (1:50). (**B**) Chemical structures and reaction scheme of phospholipase C (PLC) mediated PIP_2_ hydrolysis. (**C**) Overlay of selected regions of 2D ^1^H-^15^N NMR spectra of αSyn bound to PC-PIP_2_ vesicles (1:50) before (green) and after PLC hydrolysis (dark grey). N-terminal residues 1-10 are highlighted. Corresponding residue-resolved signal attenuation profiles (I/I_0_) of free versus PC-PIP_2_ vesicle-bound αSyn before (green) and after PLC hydrolysis (dark grey). Positions of C-terminal αSyn proline residues without peptide amide resonances are shown in the three-letter amino acid code.

**Figure 2 - Supplement 4:**
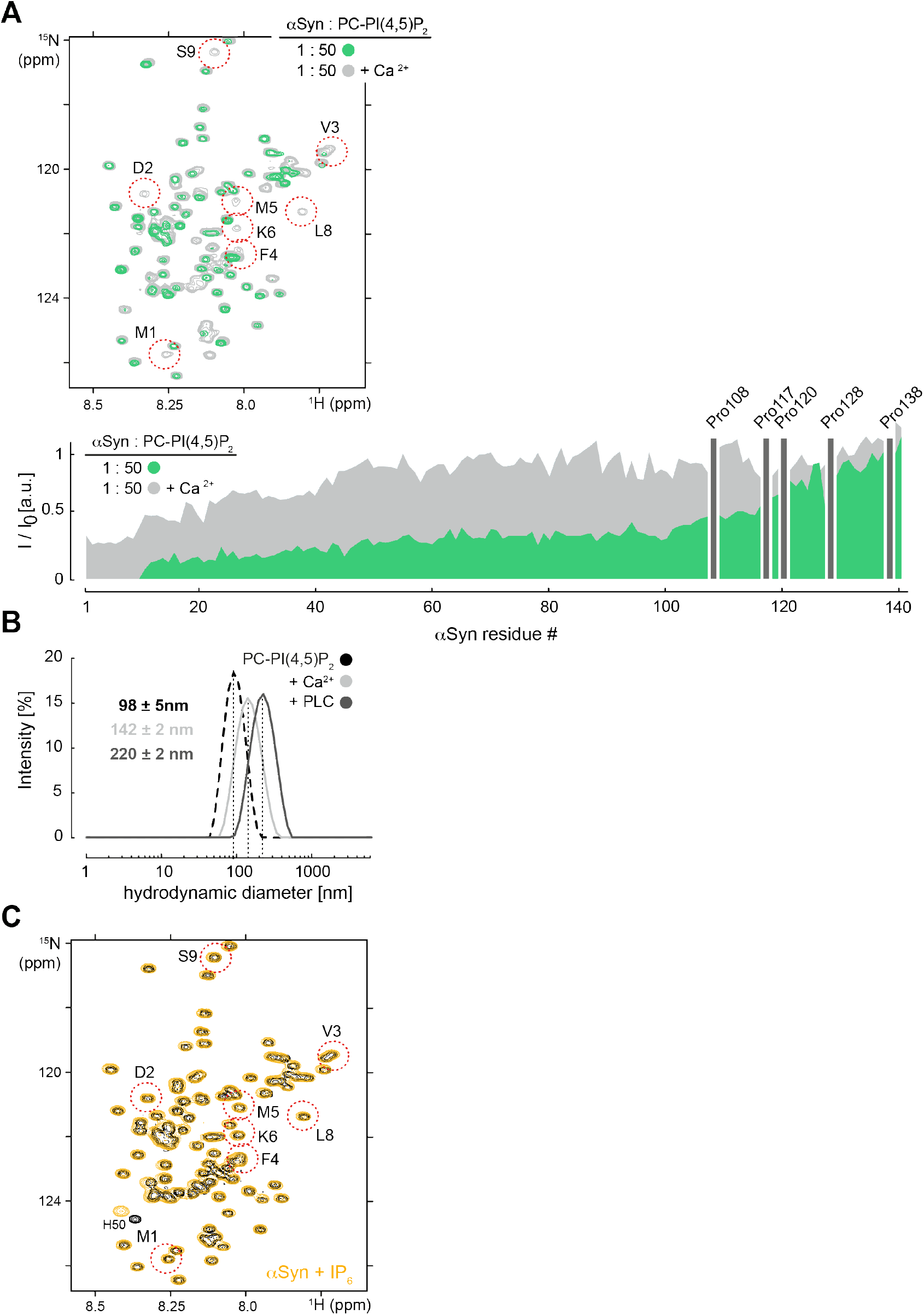
(**A**) Overlay of 2D ^1^H-^15^N NMR spectra of PC-PIP_2_ vesicle-bound αSyn in the absence (green) and presence of Ca^2+^ (light grey). Residue-resolved NMR signal intensities ratios (I/I_0_) of free αSyn versus PC-PIP_2_ vesicle-bound αSyn (I) with (green) and without Ca^2+^ (light grey). Positions of C-terminal αSyn proline residues without peptide amide resonances are shown in the three-letter amino acid code. (**B**) Hydrodynamic diameters of free and αSyn-bound PC-PIP_2_ vesicles (1:50) in the presence of Ca^2+^ (light grey) or PLC (dark grey) by DLS experiments. Errors were calculated based on measurements of three independent replicate samples. (**C**) Overlay of 2D ^1^H-^15^N NMR spectrum of αSyn (black) in presence of a 4-fold molar excess of free IP_6_ (orange). N-terminal residues 1-10 are highlighted. Slight changes in sample pH towards more acidic value are indicated by the chemical shift change of αSyn H50.

**Figure 3 - Supplement 1:**
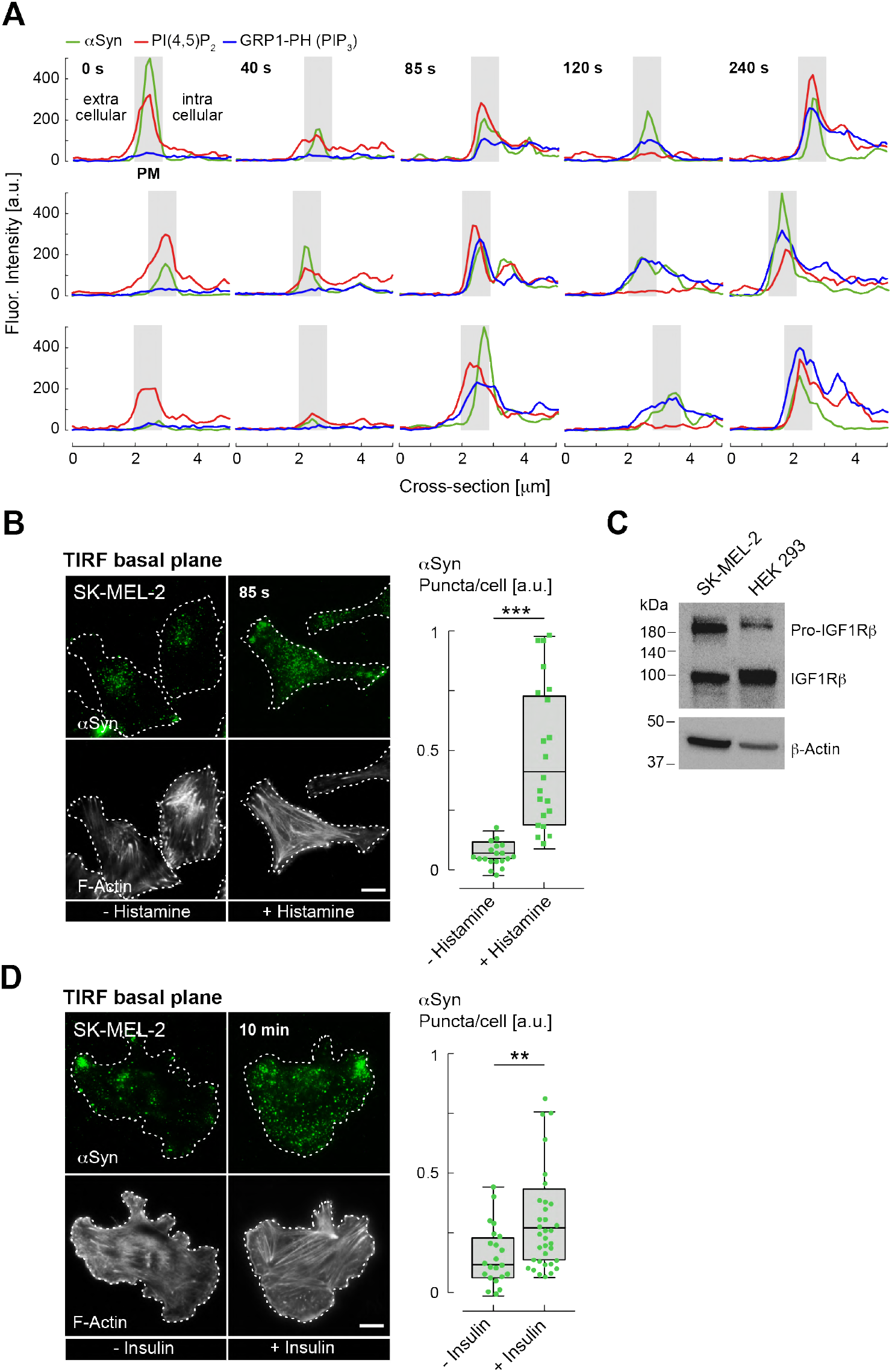
(**A**) Spatially-resolved fluorescence intensity profiles of αSyn (green), PIP_2_ (red) and GRP1-PH GFP/PIP_3_ (blue) signals at the PM at indicated time points following histamine stimulation. Individual traces span extracellular and intracellular portions of analyzed cells. PM regions are indicated by grey boxes. **(B)** and **(D)** Immunofluorescence localization of endogenous αSyn in SK-MEL-2 cells by TIRF-microscopy, counterstained with Phalloidin for F-Actin to mark cell boundaries. Cells were stimulated with histamine (B) or insulin (D). Quantification of αSyn signals at basal PM regions with and without stimulation shown on the right. Box plots represent data points collected from n= ~120 cells combined from three independent replicate experiments. Significance based on Student’s *t* tests as (**)P < 0.01; (***)P < 0.001. Scale bars are 10 μm. **(C)** Western blot of SK-MEL-2 and HEK 293 cell lysates showing the presence of endogenous insulin like growth factor receptor β (IGF-Rβ). β-Actin serves as loading-control.

